# A progressive reduction in basal phosphorylation of glutamate receptors along postweaning development accompanies maturation of hippocampal theta-burst-induced LTP

**DOI:** 10.64898/2026.07.30.741885

**Authors:** M Bento, V Guerreiro-Pinto, M Gil, NC Rodrigues, D Cunha-Reis

**Affiliations:** Biosystems and Integrative Sciences Institute (BioISI), Universidade de Lisboa, Campo Grande, 1749-016 Lisboa, Portugal; Departamento de Biologia (DBio), Faculdade de Ciências, Universidade de Lisboa, Campo Grande, 1749-016 Lisboa, Portugal; Unidade de Neurociências, Instituto de Medicina Molecular, Faculdade de Medicina, Universidade de Lisboa, Av. Prof. Egas Moniz, 1649-028 Lisboa, Portugal

**Keywords:** theta-burst, hippocampus, AMPA Rs, NMDA Rs, CaMKII, PKC/PKM, synaptophysin, LTP, phosphorylation

## Abstract

Tackling developmental changes in long-term potentiation (LTP) expression during adolescence is crucial to understand disease susceptibility in neurodevelopmental disorders, epileptogenesis or drug abuse. This paper investigated the alterations in the ‘resting’ phosphorylation state of synaptic proteins essential for synaptic transmission and hippocampal LTP expression from weaning (3 weeks) to adulthood (12 weeks). Synaptic AMPA GluA1 and GluA2 subunit levels increased, yet to different extents, as the GluA1/GluA2 ratio also increased. Conversely, GluA1 phosphorylation in both Ser831 and Ser845 progressively decreased from weaning to adulthood, suggesting an enhanced availability of these sites for activity-dependent phosphorylation. In accordance, LTP induced by different theta-burst stimulation (TBS) intensities in the CA1 area of rat hippocampal slices enhanced gradually in this developmental period. In contrast, basal phosphorylation of GluN1, GluN2B and CaMKII increased during adolescence. Altogether, these findings suggest that alterations in AMPA receptor subunit composition and basal phosphorylation are crucial for the maturation of hippocampal LTP during adolescence, while a mild enhancement in synaptic CaMKII levels may further provide the necessary structural support to LTP expression and stability. Given the reported involvement of GluA1 phosphorylation changes in epileptogenesis and neurodevelopmental disorders, these findings provide important insights into hippocampal synaptic plasticity in normal and altered brain development and epileptogenesis.

## Introduction

Brain circuit development occurs from embryonic stages to late adolescence [1,2]. Recent and accumulated evidence suggests that distinct brain regions show differential patterns of developmental plasticity from childhood to adolescence, a developmental period associated with the development of higher-order cognitive function[2,3] and during which there is a large susceptibility to the emergence intellectual and social disabilities in response to environmental factors such as social stress or drug abuse [2,4,5]. One of the hallmarks of this developmental period is a progressive maturation of synaptic plasticity towards adulthood, allowing for circuit stabilisation while maintaining adaptability [1,6,7].

Long-term potentiation (LTP), a form of activity dependent synaptic plasticity, is widely accepted as a cellular model for synaptic adaptation processes involved in hippocampal memory storage [8,9]. Theta-burst stimulation (TBS), commonly used to trigger LTP, consists of a sequence of electrical stimuli that mimics CA3 and CA1 pyramidal complex-spike cell discharges concurrent with the hippocampal theta rhythm (3-7Hz), an EEG pattern observed during hippocampal spatial memory storage, believed to act as a ‘tag’ for short term memory processing [9,10]. Importantly, TBS acts by suppressing feedforward inhibition through a priming burst or single pulse, activating GABA_B_ autoreceptors, unleashing GABA release from feedforward interneurons [11] and increases synaptic GABA availability. This allows sufficient depolarization to activate NMDA receptors [12,13] and trigger an early-LTP [8].

TBS-induced LTP in the hippocampus relies on the activity-dependent activation of NMDA receptors triggering Ca^2+^ influx into the postsynaptic compartment, the subsequent activation and auto-phosphorylation of Ca^2+^/calmodulin-dependent protein kinase II (CaMKII) [14,15] leading to the phosphorylation of AMPA receptor GluA1 subunits, enhancing the channel conductance and leading to its recruitment to the active zone of the postsynaptic membrane (early-LTP) [16–20]. Nevertheless, recent evidence also implicates CaMKII in LTP induction through binding to the GluN2B subunit of NMDA receptors through a phosphorylation-dependent structural function [21,22]. LTP stability and long-lasting expression involves the subsequent activation of multiple intracellular signalling cascades [8,23], many of which impact the activity/phosphorylation of AMPA and NMDA receptor subunits, enzymes that regulate glutamate channel activity or yet impact LTP expression and maintenance, such as the continuously active atypical protein kinase C ζ (PKCζ), PKMζ, that is critical for enduring maintenance of wild-type LTP [24,25] or the ubiquitous protein kinase A (PKA), that is implicated in LTP maintenance, late-LTP and is crucial for memory capacity [17,26].

In the hippocampus, synaptic plasticity is strongly influenced by the activity of hippocampal GABAergic interneurons that impact not only the balance between excitation and inhibition but still a complex network of circuits regulating feedforward, feedback, and disinhibitory mechanisms [27,28]. In fact, Ca^2+^-dependent input selectivity and precision of LTP induction is majorly influenced by synaptic inhibition at pyramidal cell dendrites [29]. Despite recent advances [30], the long lasting plasticity responses of interneurons to mild TBS remain largely unknown. We recently reported that LTP induced by TBS undergoes postweaning developmental changes [6] that are prospectively dependent on maturation of GABAergic circuits [7]. Furthermore, mounting evidence suggests that interneurons mediate normal circuit development through regulation synaptic plasticity at critical developmental periods, like adolescence, while shaping the formation of sensory maps [27,31]. This in turn influences overall glutamatergic synaptic activity and likely impacts the basal levels of AMPA and NMDA subunit phosphorylation that govern hippocampal LTP induction and expression [18,32].

In this paper we investigated the postweaning developmental changes in the basal phosphorylation levels of enzymes and channels relevant for synaptic plasticity including AMPA and NMDA receptors and CaMKII and its link to CA1 hippocampal LTP induced by *mild* to *strong* TBS. A preliminary report of the results described was previously published in conference proceedings [33].

## Material and Methods

The experiments were performed in *juvenile* (3 weeks old), *young adult* (6-7 weeks old) and *adult* (12-14 weeks old) male outbred Wistar rats (n=86) from Harlan Iberica or Charles River Laboratories (Barcelona, Spain) essentially as previously described [7,34] and all protocols and procedures were performed according to ARRIVE guidelines for experimental design, analysis, and their reporting. Animal housing and handing was performed in accordance with the Portuguese law (DL 113/2013) and European Community guidelines (86/609/EEC and 63/2010/CE). The animals were anesthetized with fluothane, decapitated, and the right hippocampus dissected free in ice-cold artificial cerebrospinal fluid (aCSF) composed of in mM: NaCl 124, KCl 3, NaH_2_PO_4_ 1.25, NaHCO_3_ 26, MgSO_4_ 1.5, CaCl_2_ 2, glucose 10, and gassed with a 95% O_2_ - 5% CO_2_ mixture.

### Western blot analysis of synaptic proteins

The hippocampi of 3 week-old, 6-7-week-old and 12 week-old rats were collected in sucrose solution (320mM Sucrose, 1mg/ml BSA, 10mM HEPES e 1mM EDTA, pH 7,4) containing protease and phosphatase inhibitors (MS-SAFE Protease and Phosphatase Inhibitor Cocktail, Sigma-Aldrich), homogenized with a Potter-Elvejham apparatus and hippocampal synaptosomes were isolated as described [7,35]. Hippocampal tissue homogenates were first centrifuged at 1500 g for 10 min at 4° C and the supernatant was then centrifuged at 14 000 g for 12 min at 4° C. The pellet was resuspended in 3 ml of a Percoll 45% (v/v) in modified aCSF (20mM HEPES, 1mM MgCl_2_, 1.2mM NaH_2_PO_4_, 120mM NaCl; 2.7mM KCl, 1.2mM CaCl_2_, 10mM glucose, pH 7.4) also containing protease and phosphatase inhibitors. The top layer (synaptosomal fraction) obtained upon centrifugation at 14 000 g for 4 min at 4°C, was washed twice and resuspended in modified aCSF to a final concentration of 1mg/ml protein as determined by the Bradford assay. Aliquots of 10-20 µg of these hippocampal synaptosome suspensions were stored at -80°C until use.

For western blot, samples incubated at 95°C for 5 min with Laemmli buffer (125mM Tris-BASE, 4% SDS, 50% glycerol, 0,02% Bromophenol Blue, 10% β-mercaptoethanol), were run on standard 10% sodium dodecyl sulphate polyacrylamide gel electrophoresis (SDS-PAGE) then transferred to PVDF membranes (pore size 0.45 μm, GE Healthcare Life Sciences). Membranes were then incubated for 1 h with either a 3% BSA or 5% milk blocking solution at RT and later overnight at 4°C with one of the following primary antibodies: rabbit anti GluA1 (1:4000, Millipore Cat# AB1504; RRID:AB_2113602), rabbit monoclonal anti phospho-Ser845-GluA1 (1:2500, Abcam Cat# Ab76321; RRID: AB_1523688), rabbit monoclonal anti phospho-Ser-831-GluA1 (1:2000, Abcam Cat# Ab109464; RRID: AB_10862154), rabbit anti GluA2 (1:1000, Proteintech Cat# 11994-1-AP; RRID: AB_2113725), mouse monoclonal anti GluN1 (1:1000, Proteintech Cat# 67717-1-Ig; RRID: AB_2882906), rabbit anti phospho-Ser890-GluN1 (1:500, ABclonal Cat# AP0826; RRID: AB_ 2771145), mouse monoclonal anti GluN2A (1:1000, Biolegend Cat# 832102; RRID: AB_2734597), rabbit anti GluN2B (1:1000, Cell Signalling Technology Cat# 4207; RRID: AB_1264223), mouse monoclonal anti CaMKII (1:500, Santa Cruz Biotechnology, Cat# sc-13141, RRID: AB_ 626789), mouse monoclonal anti p-CaMKII (1:500, Santa Cruz Biotechnology, Cat# sc-32289, RRID: AB_ 626786), mouse monoclonal anti PKC3 (1:500, Santa Cruz Biotechnology, Cat# sc-17781, RRID: AB_ 628148), rabbit monoclonal anti P-PKC3 (1:500, Abclonal, Cat# AP1149, RRID: AB_ 628148), rabbit anti-synaptophysin (1:7500, Synaptic Systems Cat# 101002, RRID:AB_2864013), and either mouse monoclonal anti-β-actin (1:5000, Proteintech Cat# 60008-1-Ig, RRID: AB_2289225) or rabbit polyclonal anti-α-tubulin (1:4000, Proteintech Cat# PT11224-1-AP, RRID: AB_ 2210206) primary antibodies. After washing 3x for 10 min with TBST, the membranes were incubated for 1h with anti-rabbit IgG or anti-mouse IgG secondary antibody both conjugated with horseradish peroxidase (HRP) (Proteintech) at room temperature. HRP activity was visualized by enhanced chemiluminescence with Clarity ECL Western Blotting Detection System (Bio-Rad). Intensity of the bands was evaluated with the Image J software using alpha-tubulin or beta-actin band density as a loading control. Immunostaining of the different targets was normalised to loading control band density, and differences in target protein expression from weaning to adulthood were expressed as percentage change relative to *juvenile* animals.

### LTP experiments

Hippocampal slices (400 µm thick) were cut perpendicularly to the long axis of the hippocampus with a McIlwain tissue chopper, then they were kept in a resting chamber in gassed aCSF at room temperature 22 °C – 25 °C for at least 1 h thus allowing for functional and energetic recovery. Each slice was moved at a time to a recording chamber for submerged slices of 1 ml capacity (Harvard Apparatus, UK) and continuously superfused at a rate of 3 ml/min with 95% O_2_ - 5% CO_2_ gassed aCSF at 30.5°C. Electrophysiological recordings in hippocampal slices were performed by stimulating two separate sets (S1 and S2) of the Schaffer collateral/commissural fibres in the *stratum radiatum* with rectangular pulses of 0.1 ms using bipolar concentric wire electrodes. Responses were evoked alternately on the two pathways every 10 s and hence each pathway was stimulated each 20 s (0.05Hz). Evoked field excitatory post-synaptic potentials (fEPSPs, Fig. 1.A) were recorded extracellularly from CA1 *stratum radiatum* using aCSF-filled micropipettes. The chosen stimulus intensity elicited a fEPSP of 650–920 mV amplitude (about 50% of the maximal response) set to a similar magnitude in both pathways, while curtailing contamination by the population spike. The averages of six consecutive fEPSP responses from each pathway were obtained, measured, graphically plotted and recorded for further analysis with a personal computer using the LTP software [36]. To determine the fEPSPs intensity we measured the slope of the initial phase of the potential. The independence of the two pathways was probed by examining paired-pulse facilitation (PPF) across both pathways prior to concluding experiments. PPF was elicited through sequential stimulation of the S1 and S2 Schaffer pathways with a 50 ms interval between pulses. The ratio between the fEPSP slopes elicited by the second (P2) and the first (P1) stimuli, i.e., P2/P1, was used to quantify synaptic facilitation. In all experiments considered for this paper less than 10% facilitation was observed.

**Figure 1.**
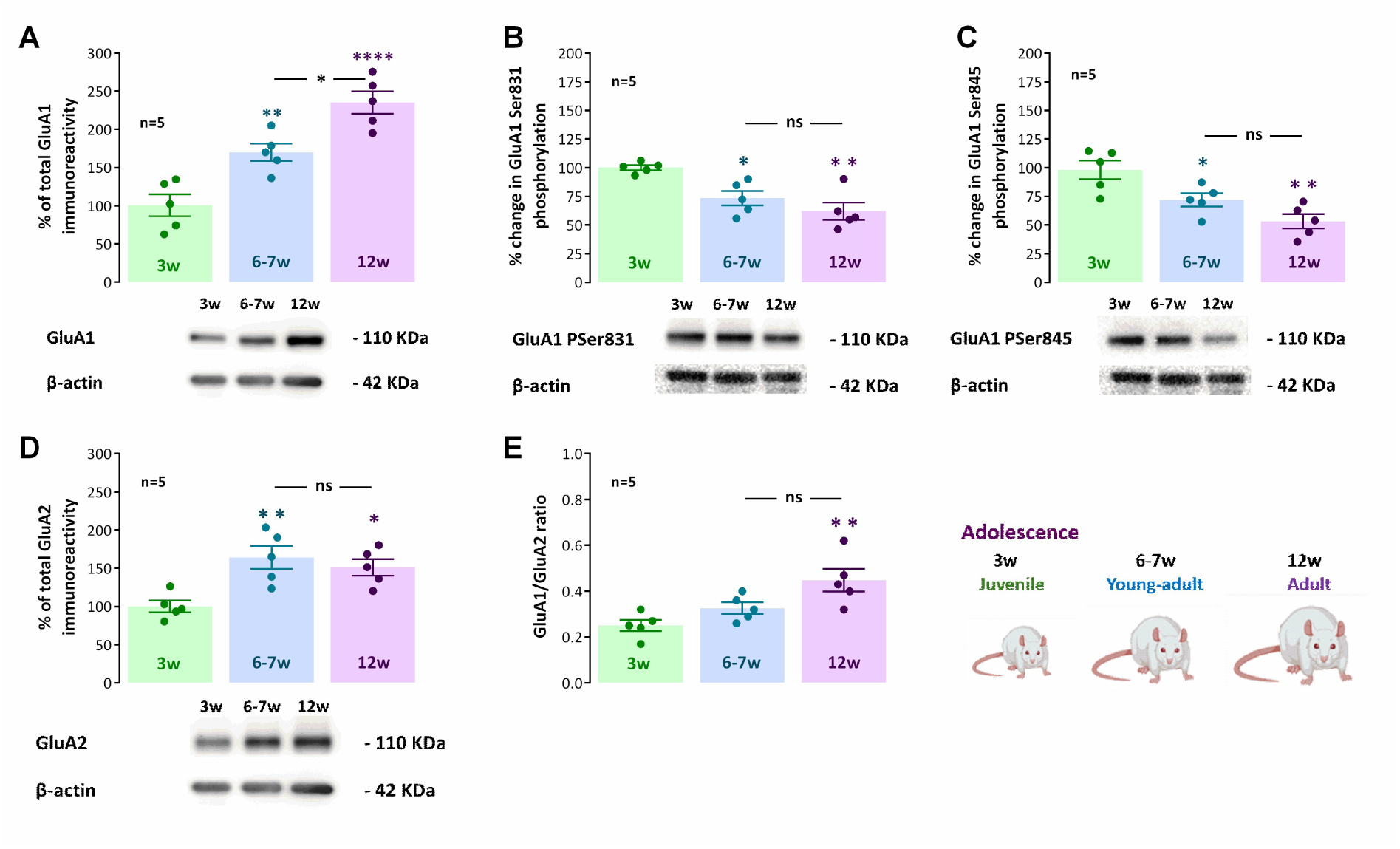
Influence of postweaning development on hippocampal GluA1 AMPA receptor phosphorylation on Ser_845_ and Ser_831_ and AMPA subunit composition. The average change in total GluA1 (A.) and GluA2 (D.) immunoreactivity (top) and western blot immunodetection of AMPA total GluA1 (A.) and GluA2 (D.) subunits (bottom) obtained in one individual experiment for *juvenile* (3 weeks), *young adult* (6-7 weeks) and *adult* (12 weeks) rats is depicted. Also shown is the % change in AMPA GluA1 Ser_831_ (B.) and Ser_845_ (C.) phosphorylation (top) and one original western blot immunodetection (bottom) of AMPA GluA1 Ser_845_ (B.) and Ser_831_ (C.) phosphorylated forms obtained in one individual experiment (bottom) for *juvenile* (3 weeks), *young adult* (6-7 weeks) and *adult* (12 weeks) rats. E. relative change of GluA1 vs. GluA2 along post-weaning development as given by the GluA1/GluA2 ratio for *juvenile*, *young adult* and *adult* rats. Values are mean ± *S.E.M.* of five independent experiments performed in duplicate. 100% - averaged GluA1 or GluA2 immunoreactivity, GluA1 phosphorylation or GluA1/GluA2 ratio obtained for *juvenile* rats. * P < 0.05, ** P < 0.01, **** P < 0.0001, *ns* - P > 0.05 (One-way *ANOVA*, using Holm-Sidak’s correction for multiple comparisons) taking as reference protein levels, GluA1 phosphorylation or GluA1/GluA2 ratio in *juvenile* rats or as indicated.

When a stable fEPSP slope baseline was observed for at least 20 min upon basal stimulation, LTP was induced either by *mild TBS* (five trains of 100 Hz, 4 stimuli, separated by 200 ms), *moderate TBS* (fifteen trains of 100 Hz, 4 stimuli, separated by 200 ms) or *strong TBS* stimulation, consisting of three moderate TBS stimulation trains delivered with a 6 min interval. The intensity of the stimulus was not changed during these protocols. LTP was taken as the % change in the average slope of fEPSPs observed from 50 to 60 min after the induction protocol, in relation to one measured during the 12 min that preceded TBS. Control and test conditions were tested in independent pathways in the same slice. In all experiments, S1 always refers to the pathway (left or right, randomly assigned) to which the TBS was applied. The control pathway did not receive TBS stimulation and served to monitor eventual decline in fEPSP slope due to recording electrode movement or loss of viability of the slice.

### Experimental design and Statistics

In this study, a total of 77 animals were used in electrophysiological experiments among juvenile (n=20), young-adult (n=26) and adult (n=31). Since most of the electrophysiological recordings were performed as control experiments for other studies, and we used all the available recordings performed in the same conditions, we did not perform a priori calculations to determine sample size per condition. For WB experiments hippocampi from a total of 19 animals were used (10 of these were the contralateral hippocampus of animals used for electrophysiological studies) including juvenile (n=6), young-adult (n=7) and adult (n=6).

Protein levels and LTP values are depicted as the mean ± S.E.M of *n* experiments. The *n* denotes the number of animals in all figures. We used one slice per animal and condition in all electrophysiological experiments. Protein levels in synaptosomes obtained from one individual animal result from western blot experiments performed in duplicate, unless otherwise stated.

The significance of the differences between the means was calculated using One-way ANOVA with Tukey’s post-hoc test (when F was significant) when comparing LTP expression or the levels of synaptic proteins for different age groups. All statistical analysis was performed using GraphPad Prism 6.01. P values of 0.05 or less were considered to represent statistically significant differences. Level of significance within the figures is denoted as follows: *p < 0.05; **p < 0.01; ***p < 0.001; ****p < 0.0001.

## Results

To elucidate how changes in AMPA and NMDA receptor subunits, CaMKII and PKM3 enzyme levels and respective phosphorylation status during postweaning development were impacting hippocampal synaptic transmission and synaptic plasticity we performed western blot using hippocampal synaptosomes obtained from juvenile, young-adult and adult rats. This is a preparation that is composed mainly of presynaptic nerve terminals, but often maintaining attached part of the postsynaptic density [37].

The protein levels of AMPA GluA1 and GluA2 subunits displayed a postweaning developmental enhancement in hippocampal synaptosomes (Fig. 1, A. and D., n=5, P<0.05), that was more pronounced for GluA1 subunits, for which the protein levels were nearly 2.5-fold higher in adult (12-week-old) than in juvenile (3-week-old, post-weaning) rats. The % increase in GluA1 levels vs juvenile animals was 170.0±13.8% (n=5) for young adults and 235.2±14.7% (n=5) for adults (Fig. 1.A). Synaptosomal GluA2 levels increased mostly within the first 3-4 weeks postweaning and then kept stable levels until adulthood with the % increase in GluA2 levels reaching 164.2±15.0% (n=5) for young adults and 151.3±10.8% (n=5) for adults (Fig. 1.D) when compared to juvenile animals (100%). Consequently, the GluA1/GluA2 ratio was enhanced in adult animals (0.448±0.050, n=5, P<0.05) when compared to juvenile animals (0.250±0.024, n=5), which suggests an enhancement in Ca^2+^-permeable AMPA receptors along post-weaning development.

In order to understand the interplay between these findings, the role of AMPA GluA1 subunits in synaptic plasticity and the role of AMPA GluA1 phosphorylation by key enzymes regulating hippocampal synaptic plasticity in this developmental period we also investigated AMPA GluA1 basal phosphorylation at Ser831, a major target of CaMKII and PKCζ, and AMPA GluA1 basal phosphorylation at Ser_845_, a major target of PKA. We observed that both phosphorylation at Ser831 and Ser845 decreased along postweaning development (Fig. 1.B and C) and that this decrease was more pronounced for Ser_845_ phosphorylation, that in *adult* animals reached around half the levels observed in *juvenile* rats (% decrease 53.2±6.3%, n=5, P<0.05), while Ser_831_ phosphorylation in adults decreased to 62.0±7.5% (n=5, P<0.05) the levels observed in *juvenile* rats.

Upon LTP induction, CaMKII is swiftly activated by post-synaptic Ca^2+^ influx that also triggers its auto-phosphorylation [14,15]. To understand if alterations in basal CaMKII levels or basal CaMKII phosphorylation at Thr286, reflecting its basal activation state, are involved in the above-described changes in GluA1 Ser831 phosphorylation, we studied its variation along post-weaning development. CaMKII increased mildly from juvenile to young-adults but was markedly enhanced when the animals reach adulthood (% increase 155.9±24.0%, n=5, P<0.05, Fig. 2.A). In contrast, phosphorylation of CaMKII was increased by 50.7±10.2% (n=5, P<0.05, Fig. 2.B) in young-adult rats and further increased by 96.0±19.9% (n=5, P<0.05, Fig. 2.B) in adult rats. This suggests that an overall developmental enhancement in CaMKII phosphorylation during adolescence precedes an enhancement in CaMKII expression.

**Figure 2.**
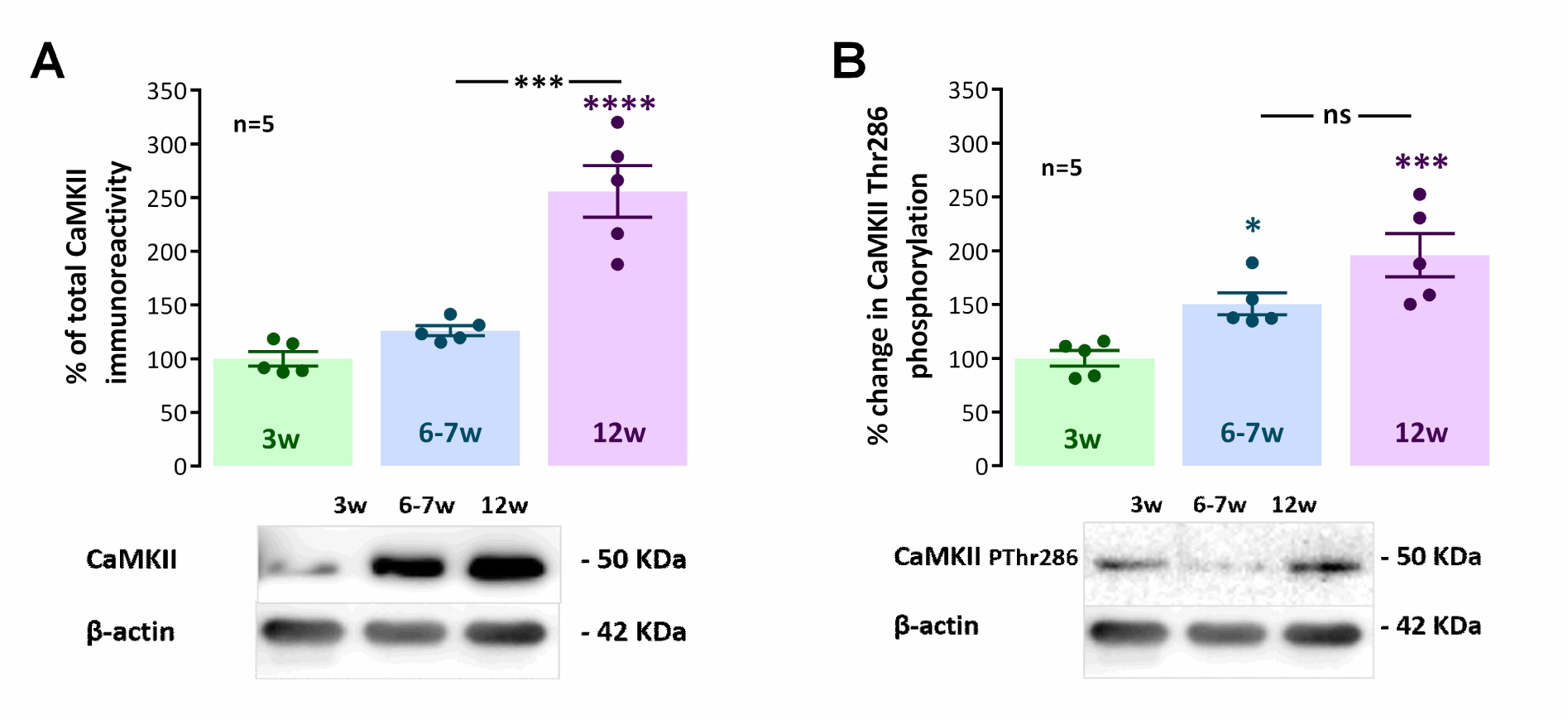
Influence of postweaning development on hippocampal CaMKII and PKC3 levels and phosphorylation. The average change in total CaMKII (**A.**) immunoreactivity (top) and original western blot immunodetection (bottom) of CaMKII obtained in one individual experiment for *juvenile*, *young adult* and *adult* rats is depicted. Also shown is the % change in CaMKII Thr_286_ phosphorylation (top) and one original western blot immunodetection (bottom) of CaMKII Thr_286_ phosphorylated form obtained in one individual experiment (bottom) for *juvenile* (3 weeks), *young adult* (6-7 weeks) and *adult* (12 weeks) rats. Values are mean ± *S.E.M.* of five independent experiments performed in duplicate. 100% - averaged CaMKII or CaMKII phosphorylation obtained for *juvenile* rats. **\*** P>0.05, **\*\*\*** P < 0.001, **\*\*\*\*** P < 0.0001, ***ns*** - P > 0.05. (One-Way *ANOVA*, using Holm-Sidak’s correction for multiple comparisons) taking as reference protein levels or CaMKII phosphorylation in *juvenile* rats or as indicated.

Developmental changes in NMDA receptor GluN1 and GluN2B subunits were never investigated during adolescence in the rat. Changes in GluN1 subunit phosphorylation at Ser890 are reported in epileptogenesis or after putative epileptogenic stimuli [38,39], known to induce a reversal of several features related to circuit maturation and control of excitability. As such we probed the changes in the levels and phosphorylation hippocampal of GluN1 subunits and its phosphorylation at Ser890 along post-weaning development. GluN1 subunit levels did not significantly change (P>0.05) in young-adult rats but were markedly enhanced by 68.0±27.1% (n=5, P<0.05, Fig. 3.A) in adult rats. GluN1 phosphorylation was strongly but transiently increased by 118.6±39.1% (n=5, P<0.05, Fig. 3.B) in young-adult rats and back to baseline levels in adult rats (n=5, P>0.05, Fig. 3.B). Again, this transient increase in GluN1 Ser890 phosphorylation appeared to precede the enhancement in hippocampal adulthood elevation in GluN1 levels.

**Figure 3.**
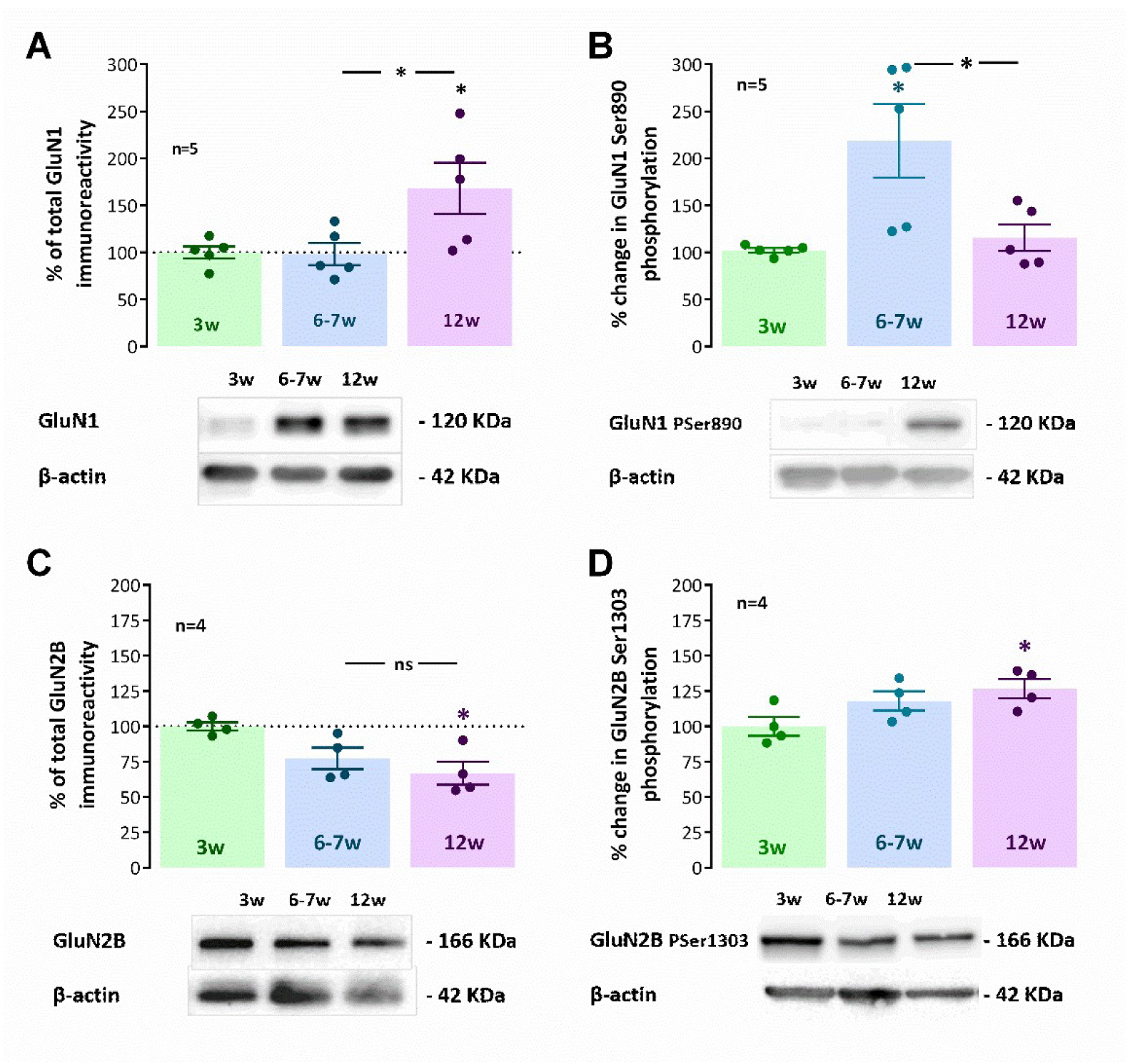
Impact of postweaning development on hippocampal levels of GluN1 and GluN2B and respective phosphorylation. The average change in total GluN1 (**A.**) and total GluN2B (**C.**) immunoreactivity (top) and original western blot immunodetection (bottom) of GluN1 (**A.**) and total GluN2B (**C.**) subunits obtained in one individual experiment for *juvenile*, *young adult* and *adult* rats is depicted. Also shown is the % change in GluN1 Ser_890_ (**B.**) and GluN2B Ser_1303_ (**D.**) phosphorylation (top) and one original western blot immunodetection (bottom) of GluN1 Ser_890_ (**B.**) and GluN2B Ser_1303_ (**D.**) phosphorylated forms obtained in one individual experiment (bottom) for *juvenile* (3 weeks), *young adult* (6-7 weeks) and *adult* (12 weeks) rats. Values are mean ± *S.E.M.* of four to five independent experiments performed in duplicate. 100% - averaged GluN1/GluN2B immunoreactivity or GluN1/GluN2B phosphorylation obtained for *juvenile* rats. **\*** P < 0.05, ***ns*** - P > 0.05 (One-Way *ANOVA*, using Holm-Sidak’s correction for multiple comparisons) taking as reference protein levels or GluN1/GluN2B phosphorylation in *juvenile* rats or as indicated.

GluN2B subunits are a major target of CaMKIIα that regulates overall NMDA2B synaptic availability and can contribute to the overall enhancement of synaptic strength during development. As such, we probed for GluN2B levels and GluN2B phosphorylation at Ser1303, that is targeted by CaMKII [40]. Hippocampal GluN2B subunit levels underwent a mild progressive decline along adolescent development reaching a % decrease of 66.9±6.3 (n=5, P<0.05, Fig. 3.C) in adult rats. Conversely, GluN2B phosphorylation at Ser1303 mildly and progressively increased from weaning to adulthood, when it reached a % increase of 26.6±6.8 (n=5, P<0.05, Fig. 3.C) when compared to juvenile animals.

To establish the relevance of these findings for hippocampal synaptic plasticity we investigated LTP induced by theta-burst stimulation in the CA1 area of the hippocampus. The fEPSP recordings obtained under basal stimulation conditions (40-60% of the maximal response in each slice) in hippocampal slices from *juvenile* rats (Fig. 4.A, raw data from a single experiment) showed an average slope of 0.579±0.036 mV/ms (n=11). When *mild TBS (5×4 pulses with a 200ms interval)* was applied to the Schaffer collateral pathway (S1) an LTP was induced, leading to a persistent 23.3±1.0% increase in fEPSP slope (n=11, P<0.05, Fig 4.B) observed within 50-60min after TBS. The non stimulated pathway (S2) did not show this potentiation (% change in fEPSP slope of -3.0±2.6%, n=11), showing that potentiation is pathway-specific. When delivering the same stimulus to slices from *young-adult* rats (6-7 weeks old), the resulting potentiation was larger (30.6±1.0%, n=12, Fig. 4.B) and even higher in *adult* 12-week-old rats (40.7±2.2%, n=11, Fig. 4.B).

**Figure 4.**
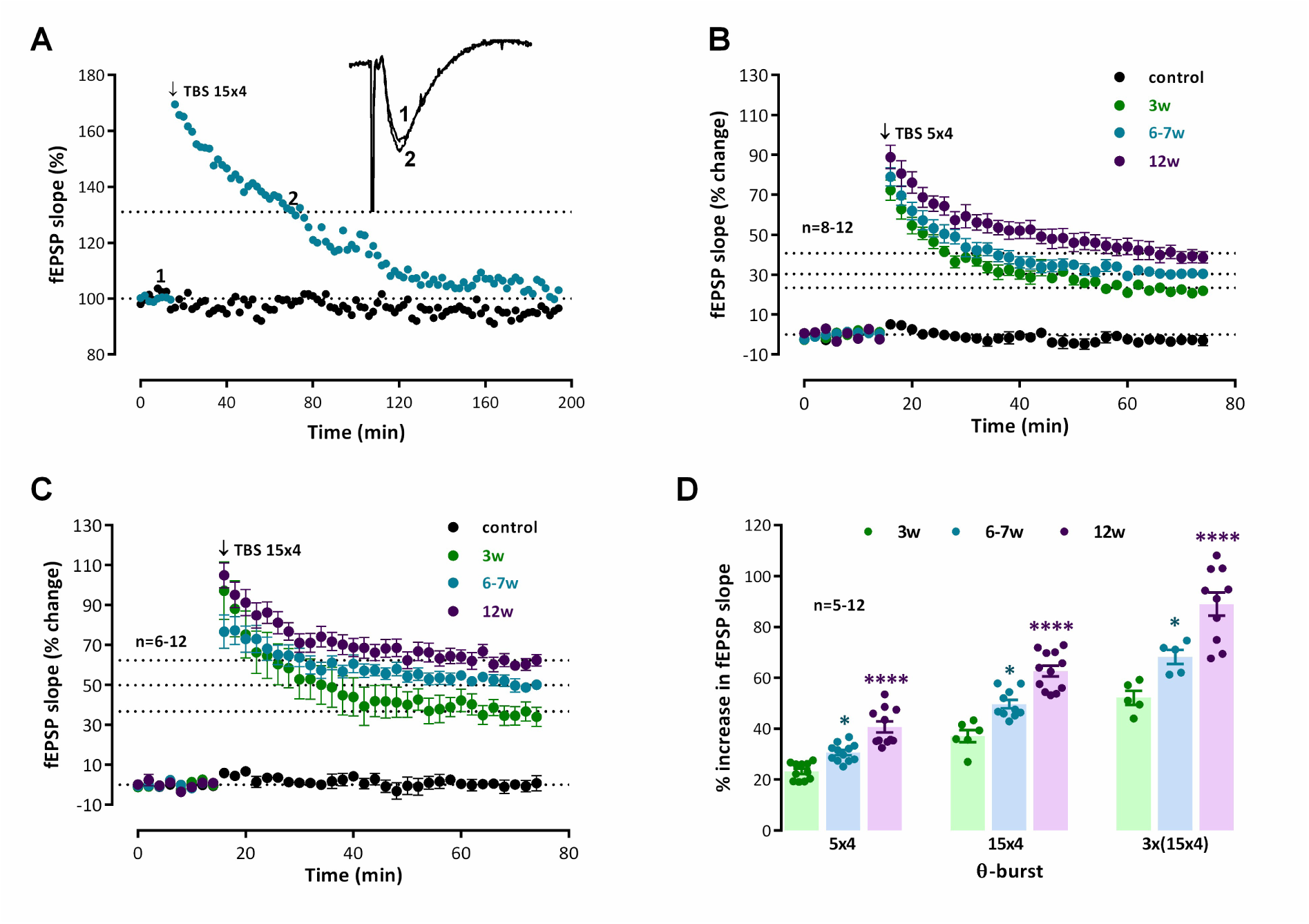
Maturation of hippocampal CA1 long-term potentiation of synaptic transmission elicited by different theta-burst stimulation intensities from weaning to adulthood. **A.** Time-course of changes in fEPSP slope caused by theta-burst stimulation (5 bursts at 5 Hz, each composed of four pulses at 100 Hz, *mild TBS (5×4)*) in a typical experiment showing both the test pathway (-•-, S2), in which *TBS (5×4)* was delivered at t=16 min and an internal control pathway (-●-, S1), obtained in the absence of TBS stimulation in the same hippocampal slice from a *juvenile* rat (3 weeks). *Inset:* Traces of fEPSPs obtained in the same experiment before (time point **1**) and 50-60 min after (time point **2**) theta burst stimulation. Traces are the average of eight consecutive responses and are composed of the stimulus artifact, the presynaptic volley and the fEPSP. Averaged time-course of changes in fEPSP slope elicited by either *mild TBS (5×4)* (shown in **B.**) or *moderate TBS (15×4)* (shown in **C.**) for *juvenile* (3 weeks, -•-), *young adult* (6-7 weeks, -•-) and *adult* (12 weeks, -•-) animals. Control and test conditions (absence and presence of TBS) were tested in independent pathways in the same slice. **D.** Average LTP magnitude obtained 50-60 min theta-burst stimulation with increasingly stronger TBS stimulation paradigms: *mild TBS (5×4), TBS (10×4), moderate TBS (15×4)* and *strong TBS 3x (15×4)* bursts separated by 6 min in the absence of added drugs in young adult rats (6-7 weeks). **D.** Average LTP magnitude obtained 50-60 min after TBS with increasingly stronger TBS stimulation paradigms: *mild TBS (5×4), moderate TBS (15×4)* and *strong TBS 3x (15×4)* in *juvenile*, *young adult* and *adult* animals. Values (**B.** - **D.**) are the mean ± S.E.M. *p < 0.05, ****p < 0.0001 (one-way *ANOVA*, Tukey’s multiple comparison test) as compared with the potentiation obtained 50-60 min after *each TBS (mild, moderate or strong TBS) for juvenile animals (3-weeks-old)*.

When delivering a stronger TBS by increasing the number of bursts to 15 (*moderate TBS*) the resulting potentiation 50-60min after TBS in *juvenile* rats was larger than when using *mild* TBS in this age group, increasing the fEPSP slope by 36.7±2.9% (n=5, P<0.05, Fig 4.C). The same *moderate TBS* pattern delivered to hippocampal slices from *young-adult* rats resulted in an even higher LTP (% increase in fEPSP slope 49.6±1.7%, n=10, P<0.05, Fig 4.C). The LTP was much higher upon *moderate TBS* delivery to *adult* rat slices and resulted in a potentiation of fEPSP slope of 62.6±2.2% (n=12, P<0.05, Fig 4.C).

By using a *strong TBS* paradigm (3x *moderate TBS* separated by 6 min) the potentiation observed in *juvenile* rats was higher (52.2±2.7%, n=5, P<0.05, Fig 4.D) than the ones observed with *mild* and *moderate TBS*. The *strong TBS* paradigm applied to hippocampal slices from *young-adult* and *adult* rats resulted in cumulatively higher LTP (% increase in fEPSP slope 68.2±2.7%, n=5 and 88.9±4.6%, n=10, respectively, P<0.05, Fig 4.D).

## **4.** Discussion

The main findings of the present work, aiming to study molecular alterations in targets relevant for synaptic plasticity in the hippocampus of male Wistar rats along adolescence from weaning (at 3 weeks) through adulthood (at 12-weeks), are that: 1) The levels of AMPA GluA1 subunits increase, while basal phosphorylation of GluA1 at Ser3831 and Ser845 decreases; 2) The AMPA receptor GluA1/GluA2 ratio increases; 3) CaMKII levels and CaMKII phosphorylation at Thr286 rises; 4) The levels of NMDA GluN1 subunits increase while GluN1 Ser890 phosphorylation increases transiently in mid-adolescence then returning to basal levels; 5) NMDA GluN2B levels mildly decrease while phosphorylation of GluN2B at Ser1303 mildly enhances and that 6) *mild to strong TBS* stimulation, a paradigm mimicking complex-spike discharges under theta-rhythm modulation that are observed during exploration and memory acquisition, induces progressively larger LTP in the CA1 area of hippocampal slices.

These results evidence that during adolescence, after the main brain structures are established, and past the developmental GABA shift [41], glutamatergic transmission undergoes still profound molecular alterations in the hippocampus (and likely in occurring in other cortical areas) that allow the establishment of fully mature synaptic plasticity, particularly LTP. By focusing on the levels and phosphorylation status of glutamatergic synaptic proteins and intracellular enzymes relevant for LTP induction and expression on a moment prior to LTP induction , this work adds to previous findings from our group showing that TBS-induced LTP undergoes post-weaning developmental maturation [6], linked to a progressive enhancement of glutamatergic and, more prominently, GABAergic synaptic contacts in the rat [7] Interestingly, this does not recapitulate in the mouse (see Kaizuka and Takumi, 2024 for review) [42].

Induction of LTP by TBS in excitatory synapses of the hippocampus involves the overthrow of feed-forward phasic inhibition, mediated by GABA_B_ autoreceptor suppression of GABA release [9,43,44], allowing for the temporal summation of excitation and a prolonged depolarization that finally leads to activation of NMDA receptors. In addition, TBS also activates cooperative postsynaptic mechanisms dependent on GABA_B_ and metabotropic glutamate receptors, leading to a sustained potentiation of fast GABA_A_ synaptic transmission in GABAergic synapses to pyramidal cells [45]. Our previous work showed that *TBS* (5-15 bursts, 4 pulses delivered at 100Hz) elicited a long-lasting potentiation of fEPSP slope in the CA1 area of rat hippocampal slices that was increasingly larger in 3-, 6-7-, and 12-week-old rats [6]. Importantly, we and others have shown that LTP induced by all these TBS paradigms is fully dependent on the activation of NMDA receptors and on CaMKII activity [6,9,12,46]. Previous studies had shown that cellular mechanisms leading to LTP expression and stability are reinforced during postnatal development [9,47,48], yet our group unequivocally showed that this reinforcement continues during adolescent development until full adulthood at 12-weeks [6,7], under these physiological stimulation patterns. Furthermore, LTP responses to increasing TBS intensities in young adult rats was linked to enhanced AMPA GluA1 Ser831 phosphorylation and Kv4.2 Ser438 phosphorylation and that the extent of TBS-induced phosphorylation was positively correlated with TBS intensity[6].

Taking all this into consideration, we hypothesized that maturation of LTP expression during adolescence, could be due to either shifts in TBS-induced Ca^2+^ dynamics, altered basal enzymatic levels/activities, altered levels of receptors and channels essential for LTP expression or altered basal phosphorylation status of these receptors and enzymes. To untangle this, we set out to investigate the alterations in the levels of AMPA receptor subunits GluA1 and GluA2, NMDA receptor subunits GluN1 and GluN2B and of CaMKII, that dictate either Ca^2+^ dynamics and Ca^2+^-dependent responses to TBS stimulation, including downstream phosphorylation processes. We also studied the basal phosphorylation status of CaMKII and GluA1, GluN1 and GluN2B, at phosphorylation targets for CaMKII, PKA and PKC, that are of relevance for the expression and modulation of hippocampal LTP. Our western blot studies were performed in hippocampal synaptosomes and thus reflect mostly alterations within synapses rather than reflecting alterations in the total number of synapses.

AMPA receptor subunit composition is known to influence LTP outcomes, as AMPA receptors are a variable tetrameric combination of GluA1-4 subunits influencing channel function [32,49]. In hippocampal synapses, GluA1/2 heterodimer pairs are the most common, although GluA2/3 heterodimer pairs are also present [32]. Notably, GluA2 containing AMPA Rs, being impermeable to Ca^2+^, limit the contribution of AMPA Rs to the postsynaptic Ca^2+^ rise (e.g., during LTP induction) [32]. In this work, we found that the levels of both AMPA GluA1 and GluA2 subunits increased progressively during adolescence (Fig. 1.A, D). This comes in parallel with a mild enhancement in the total number of glutamatergic synapses, as we previously reported [7], and suggests a progressive increase in overall glutamatergic transmission during this developmental period. Of relevance, the increase in GluA1 was more prominent than the increase in GluA2, which led to an enhancement of the GluA1/GluA2 ratio (Fig. 1.E) along post-weaning development. This suggests that, as they mature, hippocampal synapses express more AMPA receptors, yet an increasing proportion lacking GluA2 subunits, which likely leads to an enhanced capacity of AMPA receptors to contribute to the post-synaptic Ca^2+^ peak during LTP induction. Diminished levels of GluA2 containing AMPA receptors, taking as reference healthy adults, were linked to aging and pathological conditions such as brain lesions [32] and epileptogenesis [35,50], processes also linked to aberrant synaptic plasticity. Altogether this suggests that maturation of hippocampal synaptic transmission during adolescence involves both a reinforcement of glutamatergic transmission and an enhancement of the AMPA contribution for LTP induction through Ca^2+^ signalling.

Expression of TBS-induced hippocampal LTP triggers the phosphorylation of AMPA GluA1 subunits and their synaptic recruitment [6,18,51], through a mechanism dependent on CaMKII and often intracellular kinases like PKC and PKA [52]. AMPA GluA1 phosphorylation contributes to LTP expression by two mechanisms: 1) GluA1 phosphorylation at Ser831 increases the channel conductance and promotes the recruitment of AMPA receptors to the synapse [53,54] while 2) phosphorylation of GluA1 subunits at Ser845 drives AMPA receptor synaptic presence, by reducing internalization, and enhances channel open probability [32,54]. In this work we observed a progressive reduction in basal GluA1 phosphorylation at both Ser831 (CaMKII and PKC target, Fig. 1.B) and Ser845 (PKA target, Fig. 1.C). This, together with the above-mentioned increase in the number GluA1 containing AMPA receptors suggests that along adolescent hippocampal development additional phosphorylation slots are available for activity-dependent LTP expression through multiple CaMKII, PKC and PKA-dependent mechanisms, although note all relevant for expression of TBS-induced LTP.

CaMKII activity is required for expression of CA1 hippocampal TBS-induced LTP [6,51]. CaMKII activation is triggered by Ca^2+^ influx into the postsynaptic compartment, that together with calmodulin triggers the subsequent auto-phosphorylation of CaMKII [14,15], subsequently leading to the phosphorylation of AMPA receptor GluA1 subunits. In this work we report an enhancement in both basal CaMKII levels and autophosphorylation at Thr286 during postweaning development. This appears to indicate that CaMKII is more active towards adulthood than in the juvenile hippocampus. Although our work does not allow to demonstrate that this is a consequence of enhanced excitation due to increased AMPA receptor levels within glutamatergic nerve terminals, we believe this is the most likely possibility, since enhanced AMPA receptor levels will lead to larger depolarization and more GluA2-lacking Ca^2+^-permeable receptors will lead to enhanced depolarization and Ca^2+^ entry, ultimately leading to sustained and stronger CaMKII activation and likely to its stabilization through association with GluN2B [22,40,55].

NMDA receptors are the key gate to postsynaptic Ca^2+^ entry during LTP induction. NMDA receptors are large heterotetrametric ion channel proteins that in the adult hippocampus are composed of mandatory GluN1 subunits combined predominantly with GluN2A or GluN2B subunits, forming GluN1/GluN2A and GluN1/GluN2B receptor complexes, although GluN2C and GluN2D are also present, in particular early in development, but nearly absent in adulthood [55]. In this work we observed a mild and progressive decrease in hippocampal GluN2B subunits (Fig. 3.C) along adolescence. Although this is consistent with previous observations of a developmental shift in GluN2B to GluN2A containing NMDA receptors during early life [56,57], we show that this decrease continues to occur during adolescence. Moreover, we observed a progressive enhancement in basal GluN2B phosphorylation at Ser1303 (Fig. 3.D), targeted by CaMKII, that is consistent with the enhanced levels and basal autophosphorylation of CaMKII above described. GluN2B phosphorylation at Ser1303 promotes the binding of CaMKIIα to GluN2B that is crucial to the synaptic targeting and activity of CaMKII [22]. As such, these alterations are also consistent with a role for GluN2B basal phosphorylation status in the maturation of TBS-induced LTP during adolescence. In contrast to GluN2A/GluN2B, the mechanistic role of GluN1 subunits in NMDA receptor function and LTP expression is less clear, although alterations in GluN1 have been shown following ischemia or in epilepsy models. In this work we observed a delayed enhancement in GluN1 levels in late adolescence (Fig. 3.A) that was not consistently observed in all animals. This was preceded by a transient increase in GluN1 phosphorylation at Ser890 (PKC target) in mid adolescence (Fig. 3.B). These transient changes are not likely correlated with the progressive increase in TBS-induced LTP observed from weaning to adulthood, irrespective of the strength of the TBS stimulation paradigm used (Fig. 4), that was fully dependent on NMDA receptor and CaMKII activity but independent of PKA activity for all ages and TBS paradigms used [6].

In conclusion, this paper reveals that an increase in the number GluA1 containing AMPA receptors along adolescent hippocampal development together with a decrease in GluA1 phosphorylation at Ser831 and Ser845 is likely to generate additional phosphorylation slots supporting activity-dependent LTP expression through multiple CaMKII, PKC and PKA-dependent mechanisms. Nevertheless, it is not likely that all these mechanisms, especially the PKA-dependent, are involved in TBS-induced LTP in the rat hippocampus. In addition, a developmental enhancement in CaMKII levels and CaMKII Thr286 phosphorylation will likely promote stronger plasticity mechanisms and sustained CaMKII-GluN2B interactions through GluN2B Ser1303 phosphorylation. Altogether, the elucidation of these mechanisms, that likely contribute to maturation of hippocampal synaptic plasticity during adolescence into robust adult LTP induction and expression mechanisms, may prove useful to understand cognitive impairment in neurodevelopmental disorders and provide insights into effective interventions to rescue synaptic plasticity mechanisms in these diseases.

## Acknowledgements

We acknowledge the Institute of Physiology, FMUL, for animal housing facilities and Daniela Fernandes and Armando Silva-Cruz for minor technical contribution.

## Data accessibility statement

The data that support the findings of this study are available on request from the corresponding author. The data is not publicly available due to privacy or ethical restrictions.

## Ethics statement

The work in this project was approved by the Ethics committee of the Faculty of Medicine, University of Lisbon (Comissão de ética para a saúde do CHLN/FMUL). Approval was given in writing based on a detailed procedure report by the authors.

## Author contribution

***M Bento:*** formal analysis and methodology; **Guerreiro-Pinto V:** formal analysis and methodology; ***M Gil:*** formal analysis and methodology; ***NC Rodrigues:*** formal analysis and methodology; ***D Cunha-Reis:*** formal analysis and methodology, resources, supervision, funding acquisition, project administration, and writing – original draft, review and editing.

## References

1. Garduño, B.M.; Hanni, P.; Hays, C.; Cogram, P.; Insel, N.; Xu, X. How the Forebrain Transitions to Adulthood: Developmental Plasticity Markers in a Long-Lived Rodent Reveal Region Diversity and the Uniqueness of Adolescence. Front. Neurosci. 2024, 18, 1365737, doi:10.3389/FNINS.2024.1365737/BIBTEX.

2. Fuhrmann, D.; Knoll, L.J.; Blakemore, S.J. Adolescence as a Sensitive Period of Brain Development. Trends Cogn. Sci. 2015, 19, 558–566, doi:10.1016/J.TICS.2015.07.008/ASSET/1CF485FE-299C-48F3-9322-74866711F3E4/MAIN.ASSETS/GR3.SML.

3. Larsen, B.; Luna, B. Adolescence as a Neurobiological Critical Period for the Development of Higher-Order Cognition. Neurosci. Biobehav. Rev. 2018, 94, 179–195, doi:10.1016/J.NEUBIOREV.2018.09.005.

4. Davis, E.P.; Leonard, B.T.; Jirsaraie, R.J.; Keator, D.B.; Small, S.L.; Sandman, C.A.; Risbrough, V.B.; Stern, H.S.; Glynn, L.M.; Yassa, M.A.;, et al. Sex-Specific Effects of Early Life Unpredictability on Hippocampal and Amygdala Responses to Novelty in Adolescents. bioRxiv 2024, 2024.09.20.614130, doi:10.1101/2024.09.20.614130.

5. Reh, R.K.; Dias, B.G.; Nelson, C.A.; Kaufer, D.; Werker, J.F.; Kolbh, B.; Levine, J.D.; Hensch, T.K. Critical Period Regulation Acrossmultiple Timescales. Proc. Natl. Acad. Sci. U. S. A. 2020, 117, 23242–23251.

6. Rodrigues, N.C.; Silva-Cruz, A.; Caulino-Rocha, A.; Bento-Oliveira, A.; Alexandre Ribeiro, J.; Cunha-Reis, D. Hippocampal CA1 Theta Burst-Induced LTP from Weaning to Adulthood: Cellular and Molecular Mechanisms in Young Male Rats Revisited. Eur. J. Neurosci. 2021, 54, 5272–5292, doi:10.1111/ejn.15390.

7. Gil, M.; Caulino-Rocha, A.; Bento, M.; Rodrigues, N.C.; Silva-Cruz, A.; Ribeiro, J.A.; Cunha-Reis, D. Postweaning Development Influences Endogenous VPAC1 Modulation of LTP Induced by Theta-Burst Stimulation: A Link to Maturation of the Hippocampal GABAergic System. Biomolecules 2024, 14, 379, doi:10.3390/BIOM14030379/S1.

8. Bliss, T.; Collingridge, G. Persistent Memories of Long-Term Potentiation and the N -Methyl-d-Aspartate Receptor. Brain Neurosci. Adv. 2019, 3, 1–10, doi:10.1177/2398212819848213.

9. Larson, J.; Munkácsy, E. Theta-Burst LTP. Brain Res. 2015, 1621, 38–50, doi:10.1016/j.brainres.2014.10.034.

10. Vertes, R.P. Hippocampal Theta Rhythm: A Tag for Short-Term Memory. Hippocampus 2005, 15, 923–935, doi:10.1002/hipo.20118.

11. Cobb, S.R.; Manuel, N.A.; Morton, R.A.; Gill, C.H.; Collingridge, G.L.; Davies, C.H. Regulation of Depolarizing GABA(A) Receptor-Mediated Synaptic Potentials by Synaptic Activation of GABA(B) Autoreceptors in the Rat Hippocampus. Neuropharmacology 1999, 38, 1723–1732, doi:10.1016/S0028-3908(99)00158-6.

12. Larson, J.; Lynch, G. Role of N-Methyl-D-Aspartate Receptors in the Induction of Synaptic Potentiation by Burst Stimulation Patterned after the Hippocampal Theta-Rhythm. Brain Res 1988, 441, 111–118, 10.1016/0006-8993(88)91388-1.

13. Davies, C.H.; Davies, S.N.; Collingridge, G.L. Paired-Pulse Depression of Monosynaptic GABA-Mediated Inhibitory Postsynaptic Responses in Rat Hippocampus. J Physiol 1990, 424, 513–531.

14. Lisman, J.; Yasuda, R.; Raghavachari, S. Mechanisms of CaMKII Action in Long-Term Potentiation. Nat. Rev. Neurosci. 2012 133 2012, 13, 169–182, doi:10.1038/nrn3192.

15. Chang, J.Y.; Parra-Bueno, P.; Laviv, T.; Szatmari, E.M.; Lee, S.J.R.; Yasuda, R. CaMKII Autophosphorylation Is Necessary for Optimal Integration of Ca2+ Signals during LTP Induction, but Not Maintenance. Neuron 2017, 94, 800–808.e4, doi:10.1016/J.NEURON.2017.04.041.

16. Park, P.; Kang, H.; Sanderson, T.M.; Bortolotto, Z.A.; Georgiou, J.; Zhuo, M.; Kaang, B.-K.; Collingridge, G.L. The Role of Calcium-Permeable AMPARs in Long-Term Potentiation at Principal Neurons in the Rodent Hippocampus. Front. Synaptic Neurosci. 2018, 10, 42, doi:10.3389/fnsyn.2018.00042.

17. Benke, T.; Traynelis, S.F. AMPA-Type Glutamate Receptor Conductance Changes and Plasticity: Still a Lot of Noise. Neurochem. Res. 2019, 44, 539–548, doi:10.1007/s11064-018-2491-1.

18. Appleby, V.J.; Corrêa, S.A.L.; Duckworth, J.K.; Nash, J.E.; Noël, J.; Fitzjohn, S.M.; Collingridge, G.L.; Molnár, E. LTP in Hippocampal Neurons Is Associated with a CaMKII-Mediated Increase in GluA1 Surface Expression. J Neurochem 2011, 116, 530–543, doi:10.1111/j.1471-4159.2010.07133.x.

19. Derkach, V.; Barria, A.; Soderling, T.R. Ca2+/Calmodulin-Kinase II Enhances Channel Conductance of Alpha-Amino-3-Hydroxy-5-Methyl-4-Isoxazolepropionate Type Glutamate Receptors. Proc. Natl. Acad. Sci. U. S. A. 1999, 96, 3269–3274.

20. Chater, T.E.; Goda, Y. The Role of AMPA Receptors in Postsynaptic Mechanisms of Synaptic Plasticity. Front. Cell. Neurosci. 2014, 8, doi:10.3389/fncel.2014.00401.

21. Kim, K.; Saneyoshi, T.; Hosokawa, T.; Okamoto, K.; Hayashi, Y. Interplay of Enzymatic and Structural Functions of CaMKII in Long-Term Potentiation. J. Neurochem. 2016, 139, 959–972.

22. Tullis, J.E.; Larsen, M.E.; Rumian, N.L.; Freund, R.K.; Boxer, E.E.; Brown, C.N.; Coultrap, S.J.; Schulman, H.; Aoto, J.; Dell’Acqua, M.L.;, et al. LTP Induction by Structural Rather than Enzymatic Functions of CaMKII. Nat. 2023 6217977 2023, 621, 146–153, doi:10.1038/s41586-023-06465-y.

23. Baltaci, S.B.; Mogulkoc, R.; Baltaci, A.K. Molecular Mechanisms of Early and Late LTP. Neurochem. Res. 2019, 44, 281–296, doi:10.1007/s11064-018-2695-4.

24. Pastalkova, E.; Serrano, P.; Pinkhasova, D.; Wallace, E.; Fenton, A.A.; Sacktor, T.C. Storage of Spatial Information by the Maintenance Mechanism of LTP. Science (80-.). 2006, 313, 1141–1444, doi:10.1126/science.1128657.

25. Sacktor, T.C. Chapter 2 PKMζ, LTP Maintenance, and the Dynamic Molecular Biology of Memory Storage. Prog. Brain Res. 2008, 169, 27–40, doi:10.1016/S0079-6123(07)00002-7.

26. Olivito, L.; Saccone, P.; Perri, V.; Bachman, J.L.; Fragapane, P.; Mele, A.; Huganir, R.L.; De Leonibus, E. Phosphorylation of the AMPA Receptor GluA1 Subunit Regulates Memory Load Capacity. Brain Struct. Funct. 2016, 221, 591–603, doi:10.1007/S00429-014-0927-1/FIGURES/4.

27. Goff, K.M.; Goldberg, E.M. A Role for Vasoactive Intestinal Peptide Interneurons in Neurodevelopmental Disorders. Dev. Neurosci. 2021, 43, 168–180, doi:10.1159/000515264.

28. Artinian, J.; Lacaille, J.C. Disinhibition in Learning and Memory Circuits: New Vistas for Somatostatin Interneurons and Long-Term Synaptic Plasticity. Brain Res. Bull. 2018, 141, 20–26.

29. Müllner, F.E.; Wierenga, C.J.; Bonhoeffer, T. Precision of Inhibition: Dendritic Inhibition by Individual GABAergic Synapses on Hippocampal Pyramidal Cells Is Confined in Space and Time. Neuron 2015, 87, 576–589, doi:10.1016/j.neuron.2015.07.003.

30. Wiera, G.; Jabłońska, J.; Lech, A.M.; Mozrzymas, J.W. Input Specificity of NMDA-Dependent GABAergic Plasticity in the Hippocampus. Sci. Rep. 2024, 14, 1–15.

31. Stack, C.M.; Lim, M.A.; Cuasay, K.; Stone, M.M.; Seibert, K.M.; Spivak-Pohis, I.; Crawley, J.N.; Waschek, J.A.; Hill, J.M. Deficits in Social Behavior and Reversal Learning Are More Prevalent in Male Offspring of VIP Deficient Female Mice. Exp. Neurol. 2008, 211, 67–84, doi:10.1016/j.expneurol.2008.01.003.

32. Chater, T.E.; Goda, Y. The Shaping of AMPA Receptor Surface Distribution by Neuronal Activity. Front. Synaptic Neurosci. 2022, 14, 13, doi:10.3389/fnsyn.2022.833782.

33. Bento, M.; Caulino-Rocha, A.; Ribeiro, J.; Cunha-Reis, D. A Decrease in Glua1 Basal Phosphorylation Levels May Contribute To the Enhancement in Ltp Induced By Theta-Burst Stimulation Along Postweaning Development. IBRO Neurosci. Reports 2023, 15, S304, doi:10.1016/j.ibneur.2023.08.544.

34. Aidil-Carvalho, F.; Caulino-Rocha, A.; Ribeiro, J.A.; Cunha-Reis, D. Mismatch Novelty Exploration Training Shifts VPAC1 Receptor-Mediated Modulation of Hippocampal Synaptic Plasticity by Endogenous VIP in Male Rats. J. Neurosci. Res. 2024, 102, e25333, doi:10.1002/JNR.25333.

35. Carvalho-Rosa, J.D.; Rodrigues, N.C.; Silva-Cruz, A.; Vaz, S.H.; Cunha-Reis, D. Epileptiform Activity Influences Theta-Burst Induced LTP in the Adult Hippocampus: A Role for Synaptic Lipid Raft Disruption in Early Metaplasticity? Front. Cell. Neurosci. 2023, 17, 184, doi:10.3389/FNCEL.2023.1117697.

36. Anderson, W.W.; Collingridge, G.L. The LTP Program: A Data Acquisition Program for on-Line Analysis of Long-Term Potentiation and Other Synaptic Events. J Neurosci Methods 2001, 108, 71–83, doi:S0165-0270(01)00374-0 [pii].

37. Breukel, A.I.; Besselsen, E.; Ghijsen, W.E. Synaptosomes. A Model System to Study Release of Multiple Classes of Neurotransmitters. Methods Mol. Biol. 1997, 72, 33–47, doi:10.1385/0-89603-394-5:33.

38. Cheung, H.H.; Teves, L.; Wallace, M.C.; Gurd, J.W. Increased Phosphorylation of the NR1 Subunit of the NMDA Receptor Following Cerebral Ischemia. J. Neurochem. 2001, 78, 1179–1182, doi:10.1046/J.1471-4159.2001.0780051179.X;SUBPAGE:STRING:ABSTRACT;WEBSITE:WEBSITE:PERICLES;REQUESTEDJOURN AL:JOURNAL:14714159;JOURNAL:JOURNAL:14714159;WGROUP:STRING:PUBLICATION.

39. Niimura, M.; Moussa, R.; Bissoon, N.; Ikeda-Douglas, C.; Milgram, N.W.; Gurd, J.W. Changes in Phosphorylation of the NMDA Receptor in the Rat Hippocampus Induced by Status Epilepticus. J. Neurochem. 2005, 92, 1377–1385, doi:10.1111/J.1471-4159.2005.02977.X.

40. Bayer, K.U.; De Koninck, P.; Leonard, A.S.; Hell, J.W.; Schulman, H. Interaction with the NMDA Receptor Locks CaMKII in an Active Conformation. Nat. 2001 4116839 2001, 411, 801–805, doi:10.1038/35081080.

41. Ben-Ari, Y. The GABA Developmental Shift in Health and Disease. Synap. Dev. Matur. Compr. Dev. Neurosci. 2020, 277–296, doi:10.1016/B978-0-12-823672-7.00012-0.

42. Kaizuka, T.; Takumi, T. Alteration of Synaptic Protein Composition during Developmental Synapse Maturation. Eur. J. Neurosci. 2024, 59, 2894–2914, doi:10.1111/EJN.16304.

43. Pacelli, G.J.; Su, W.; Kelso, S.R. Activity-Induced Depression of Synaptic Inhibition during LTP-Inducing Patterned Stimulation. Brain Res. 1989, 486, 26–32, doi:10.1016/0006-8993(89)91273-0.

44. Grover, L.M.; Yan, C. Blockade of GABA(A) Receptors Facilitates Induction of NMDA Receptor-Independent Long-Term Potentiation. J. Neurophysiol. 1999, 81, 2814–2822, doi:10.1152/jn.1999.81.6.2814.

45. Patenaude, C.; Chapman, C.A.; Bertrand, S.; Congar, P.; Lacaille, J.C. GABAB Receptor-and Metabotropic Glutamate Receptor-Dependent Cooperative Long-Term Potentiation of Rat Hippocampal GABAA Synaptic Transmission. J. Physiol. 2003, 553, 155–167, doi:10.1113/jphysiol.2003.049015.

46. Park, P.; Sanderson, T.M.; Amici, M.; Choi, S.-L.L.; Bortolotto, Z.A.; Zhuo, M.; Kaang, B.-K.K.; Collingridge, G.L. Calcium-Permeable AMPA Receptors Mediate the Induction of the Protein Kinase A-Dependent Component of Long-Term Potentiation in the Hippocampus. J Neurosci 2016, 36, 622–631, doi:10.1523/JNEUROSCI.3625-15.2016.

47. Cao, G.; Harris, K.M. Developmental Regulation of the Late Phase of Long-Term Potentiation (L-LTP) and Metaplasticity in Hippocampal Area CA1 of the Rat. J. Neurophysiol. 2012, 107, 902–912, doi:10.1152/jn.00780.2011.

48. Kramár, E.A.; Lynch, G. Developmental and Regional Differences in the Consolidation of Long-Term Potentiation. Neuroscience 2003, 118, 387–398, doi:10.1016/S0306-4522(02)00916-8.

49. Wiltgen, B.J.; Royle, G.A.; Gray, E.E.; Abdipranoto, A.; Thangthaeng, N.; Jacobs, N.; Saab, F.; Tonegawa, S.; Heinemann, S.F.; O’Dell, T.J.;, et al. A Role for Calcium-Permeable AMPA Receptors in Synaptic Plasticity and Learning. PLoS One 2010, 5, e12818, doi:10.1371/journal.pone.0012818.

50. Serpa, A.; Bento, M.; Caulino-Rocha, A.; Pawlak, S.; Cunha-Reis, D. Opposing Reduced VPAC1 and Enhanced VPAC2 VIP Receptors in the Hippocampus of the Li2+-Pilocarpine Rat Model of Temporal Lobe Epilepsy. Neurochem. Int. 2022, 158, 105383, doi:10.1016/j.neuint.2022.105383.

51. Park, P.; Georgiou, J.; Sanderson, T.M.; Ko, K.H.; Kang, H.; Kim, J. il; Bradley, C.A.; Bortolotto, Z.A.; Zhuo, M.; Kaang, B.K.;, et al. PKA Drives an Increase in AMPA Receptor Unitary Conductance during LTP in the Hippocampus. Nat. Commun. 2021, 12, 1–15, doi:10.1038/s41467-020-20523-3.

52. Lee, H.K.; Takamiya, K.; Han, J.S.; Man, H.; Kim, C.H.; Rumbaugh, G.; Yu, S.; Ding, L.; He, C.; Petralia, R.S.;, et al. Phosphorylation of the AMPA Receptor GluR1 Subunit Is Required for Synaptic Plasticity and Retention of Spatial Memory. Cell 2003, 112, 631–643, doi:10.1016/S0092-8674(03)00122-3.

53. Barria, A.; Muller, D.; Derkach, V.; Griffith, L.C.; Soderling, T.R. Regulatory Phosphorylation of AMPA-Type Glutamate Receptors by CaM-KII during Long-Term Potentiation. Science (80-.). 1997, 276, 2042–2045, doi:10.1126/science.276.5321.2042.

54. Derkach, V.A.; Oh, M.C.; Guire, E.S.; Soderling, T.R. Regulatory Mechanisms of AMPA Receptors in Synaptic Plasticity. Nat. Rev. Neurosci. 2007, 8, 101–113.

55. Shipton, O.A.; Paulsen, O. GluN2A and GluN2B Subunit-Containing NMDA Receptors in Hippocampal Plasticity. Philos. Trans. R. Soc. B Biol. Sci. 2014, 369, doi:10.1098/RSTB.2013.0163/22481.

56. Yashiro, K.; Philpot, B.D. Regulation of NMDA Receptor Subunit Expression and Its Implications for LTD, LTP, and Metaplasticity. Neuropharmacology 2008, 55, 1081–1094, doi:10.1016/J.NEUROPHARM.2008.07.046.

57. Rodenas-Ruano, A.; Chávez, A.E.; Cossio, M.J.; Castillo, P.E.; Zukin, R.S. REST-Dependent Epigenetic Remodeling Promotes the Developmental Switch in Synaptic NMDA Receptors. Nat. Neurosci. 2012 1510 2012, 15, 1382–1390, doi:10.1038/nn.3214.

